# Machine Learning Approach for Enumeration of Circulating Cells with Diffuse *in vivo* Flow Cytometry

**DOI:** 10.64898/2026.04.21.719882

**Authors:** Mehrnoosh Emamifar, Jane Lee, Joshua Pace, Chiara Bellini, Mark Niedre

## Abstract

**Significance:** Diffuse *in vivo* flow cytometry (DiFC) is an emerging technique for enumerating rare, fluorescently-labeled circulating tumor cells (CTCs) in small animals without drawing blood samples. DiFC uses detection of transient fluorescent peaks in time-series data. Previously, we used a simple amplitude threshold-based method for identifying peak candidates, but it ignores potentially useful information in peak shape that could reduce false-positive detections from instrument noise and increase detection efficiency of lower-amplitude peaks.

**Aim:** To develop a machine learning (ML)-integrated signal processing approach for improved CTC enumeration using DiFC by distinguishing CTC peaks from artifacts.

**Approach:** We developed an ML-integrated approach that incorporates a convolutional neural network (CNN) classifier. The CNN was trained to distinguish CTC peaks from artifacts by analyzing peak amplitude and temporal shape characteristics. Performance was validated on *in-silico*, control, and CTC-bearing mouse datasets.

**Results:** The CNN classifier achieved accuracy, precision, sensitivity, and specificity exceeding 98% on test data. Compared with our previously published threshold-based approach, the ML-integrated method increased the number of correctly identified CTCs and their flow direction while reducing false detections across validation datasets.

**Conclusions:** The ML-integrated approach significantly improves DiFC CTC enumeration, enabling robustness against artifacts in noisy conditions.

## 1 Introduction

Metastasis accounts for the majority of cancer-related deaths, and one of the main pathways of spread is via the peripheral blood (PB). As such, circulating tumor cells (CTCs) may serve as an important indicator of disease progression and and response to treatment.^1, 2^ In clinical practice, conventional liquid biopsy methods for CTC enumeration involve drawing and analyzing fractionally small (7.5 mL) blood samples. However, these are known to be inaccurate (particularly for quantifying rare CTCs) and do not readily permit measurement of changes in CTC numbers over time.^3–5^ For pre-clinical studies in mice, the entire PB volume is typically drawn, which does not allow serial measurements in the same animal. To address these limitations, our lab developed “Diffuse *in vivo* Flow Cytometry” (DiFC), which enables non-invasive, continuous monitoring of fluorescently-labeled CTCs in mice.^6–9^ In the system, specially designed fiber-optic probes (Fig. 1a) are placed on the mouse skin approximately above large blood vessels in the leg or tails (Figs. 1b,c). Unlike alternative microscopy-based *in vivo* flow cytometry (IVFC) approaches,^10–13^ DiFC uses highly scattered diffuse light that allows fluorescence detection of CTCs several millimeters into tissue. As CTCs pass through the instrument field of view (FOV), a probe detects transient fluorescent events, referred to as “peaks” (Fig. 1d). We place two optical probes arranged 3 mm apart along the blood vessel. By requiring that signal peaks be sequentially detected by both probes, we can determine the direction of CTC flow and discriminate true CTC events from random motion or electronic artifacts that may also generate peak-like fluctuations in fluorescent signals (see Sec. 2.3). Our previous DiFC signal processing approach only considered any fluorescent event exceeding a fixed amplitude threshold (usually five times the local background signal standard deviation, *σ*_bg_) as a CTC peak candidate (Peak_cand_). However, our new approach considers both the amplitude and shape of peaks. True CTC peaks exhibit a characteristic waveform governed by the spatial sensitivity function of the DiFC probe, as shown by the Jacobian profile computed via Monte Carlo simulation in Fig. 1e.^14^ The sensitivity profile as a function of time was derived assuming an average blood flow speed of 100 mm/s in anesthetized mice.^6^ As such, reliance on a simple amplitude-base thresholding strategy neglects informative features encoded in the temporal shape of the signal, which may lead to false positives. Because CTCs are exceedingly rare (as low as 1 CTC per mL of PB ^15, 16^), false-positive signals are highly problematic. Mitigation of spurious detections from random signal noise required use of sufficiently high detection thresholds to minimize false positives, at the expense of potentially missing true, lower signal-to-noise ratio (SNR) peaks that fell below the threshold. Hence, an improved signal processing method that incorporates information contained in the signal shape is advantageous both for reducing false positive detections and for improving true positive counts.

**Fig 1.**
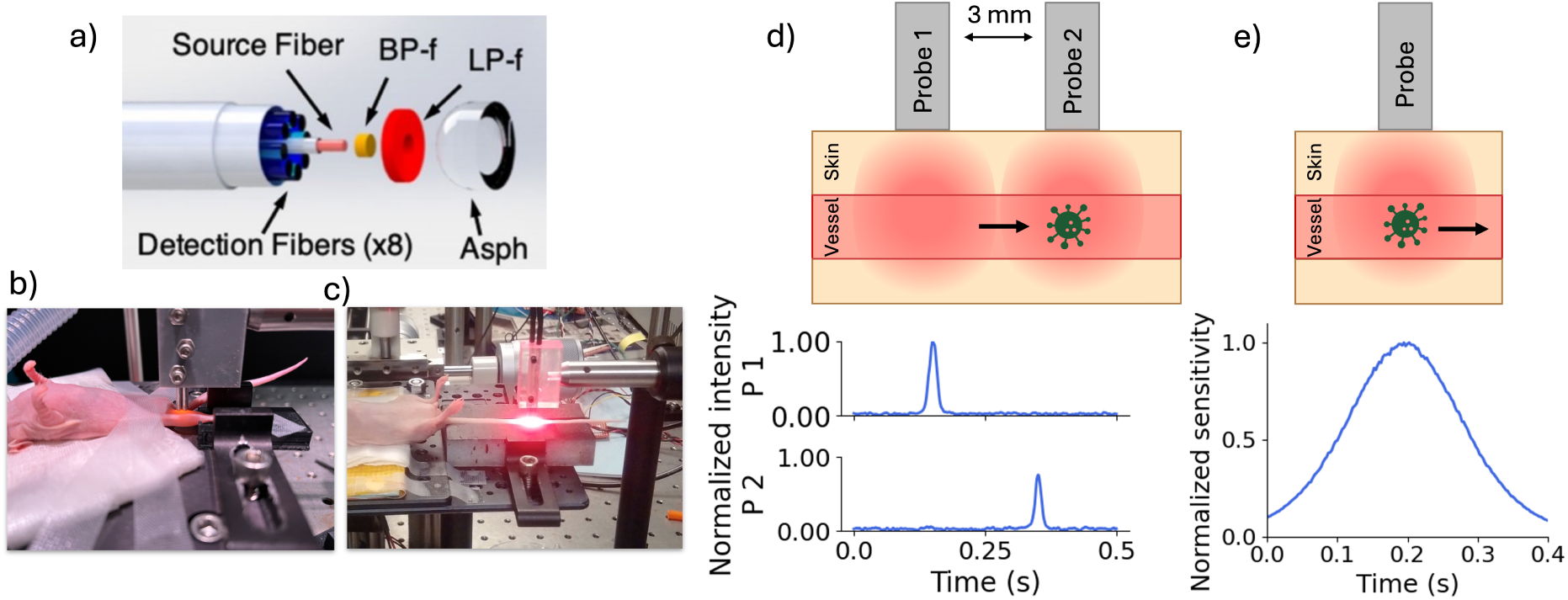
(a) Diagram of a DiFC probe, reproduced with permission from Ref. 6. We normally perform DiFC on the (b) hindlimb or (c) tail of mice. (d) Two DiFC probes are placed approximately along the blood vessel, which permits detection of fluorescently-labeled cells with a time-delay corresponding to the cell speed. (e) Peaks have a characteristic shape which is defined by the sensitivity function of the DIFC probe. An example peak shape (computed with a Monte Carlo simulation^14^) for 0.75 mm depth cells is shown.

To address this need, we present a machine learning (ML)-integrated approach that more reliably distinguishes true signal peaks from artifacts, particularly in noisy data sets.^17^ Convolutional neural networks (CNNs) are a class of ML models well suited for this task because they exhibit strong capability in extracting local features from time-series data^18^ and have been successfully applied in related signal-processing applications.^19–21^ This method is less sensitive to the choice of detection threshold and allows recovery of a greater number of true peak signals while maintaining robustness against false positives.

## 2 Methods

### 2.1 Overview of the DiFC Signal Processing Problem

The general DiFC signal processing workflow for both our previous threshold-based and the ML-integrated approach are shown in Fig. 2. During DiFC measurements, the fiber probes were placed on the skin surface approximately above a blood vessel.^6–9, 22^ Fluorescence data were collected continuously from both fiber-optic probes at a sampling rate of 2,000 samples per second. Fluorescence measurements included background signals (tissue autofluorescence), transient peaks from fluorescently-labeled CTCs, and false positive signals from animal motion (breathing, twitching) and instrument electronic noise. The overall goal of the signal processing algorithm was therefore to distinguish true CTC peaks from background and false positive signals as accurately as possible. In our previous threshold-based approach we first used signal amplitude to identify peak candidates, then a directional matching algorithm was used to reject false positives by enforcing sequential two-probe detection and excluding coincident signals as measurement noise (Fig. 2a).^6, 7^ However, by relying solely on amplitude, this approach ignored information embedded within the signal shape to aid discrimination of true peaks from false positive signals. To address this limitation, our ML-integrated approach employed a CNN classifier that leveraged both the amplitude and shape of peak candidates to reject artifacts and retain potential CTC peaks (Fig. 2b).

**Fig 2.**
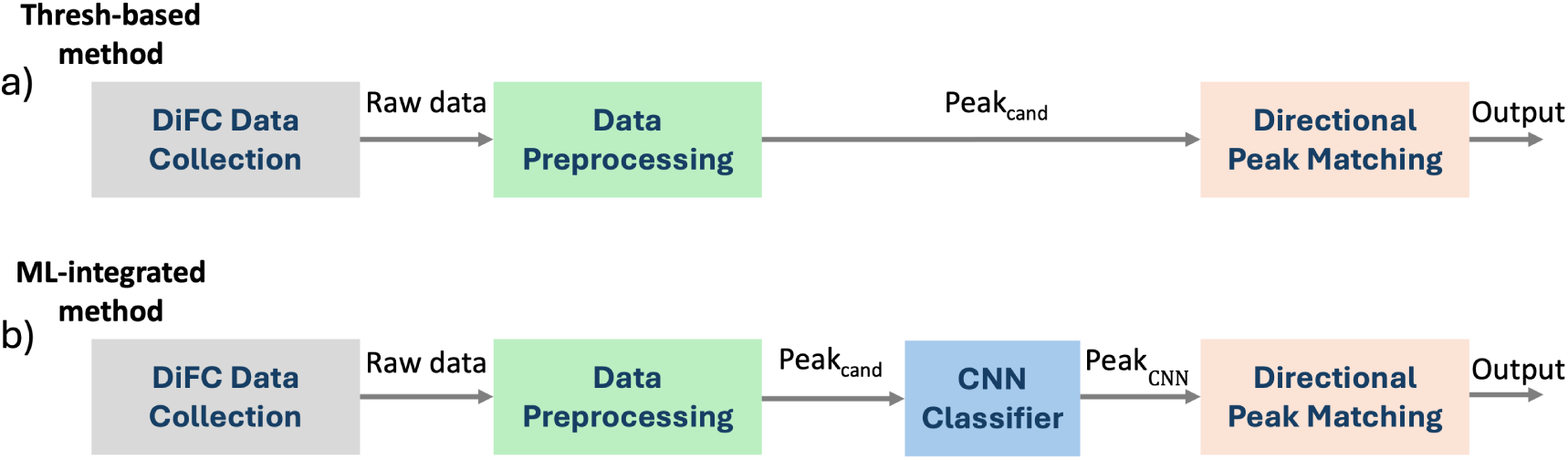
Overview of the signal processing workflow. (a) In the threshold-based approach, after pre-processing of raw data, all detected peak candidates (Peak_cand_) were processed with the directional peak matching algorithm. (b) In the ML-integrated approach, peak candidates (Peak_cand_) were first analyzed by a CNN classifier and only candidates classified as peaks (Peak_CNN_) were processed by the directional peak matching algorithm.

### 2.2 Machine Learning Model

#### 2.2.1 Signal Pre-Processing and Initial Identification of Peak Candidates

We first applied a pre-processing algorithm to the raw data acquired from both optical fibers (Fig. 3a). First, the background autofluorescence was removed from the signal by applying a sliding median filter with a window length of 2.5 s (5,000 samples) and subtracting it from the raw signal. Next, a 0.003 s (6 samples) smoothing operation was applied to the signal with a moving average filter. Peak candidates in the signal were then identified using the “find peaks” function from the “scipy.signal” module in Python, which we defined as transient signals with amplitudes exceeding a fixed multiple of the standard deviation of the local signal (4*σ*_bg_). The temporal locations of these peak candidates were recorded for further analysis. The data were normalized to [0, 1] using the minimum and maximum values of each scan to facilitate uniform processing and prevent scale-related biases. Around each detected peak candidate, a 0.4 s window (corresponding to 800 samples) was extracted from both probes and used as an input for our CNN classifier (see Sec. 2.2.2). The 0.4 s window was chosen based on analysis of peak candidate width, which consistently fit within this temporal range.

**Fig 3.**
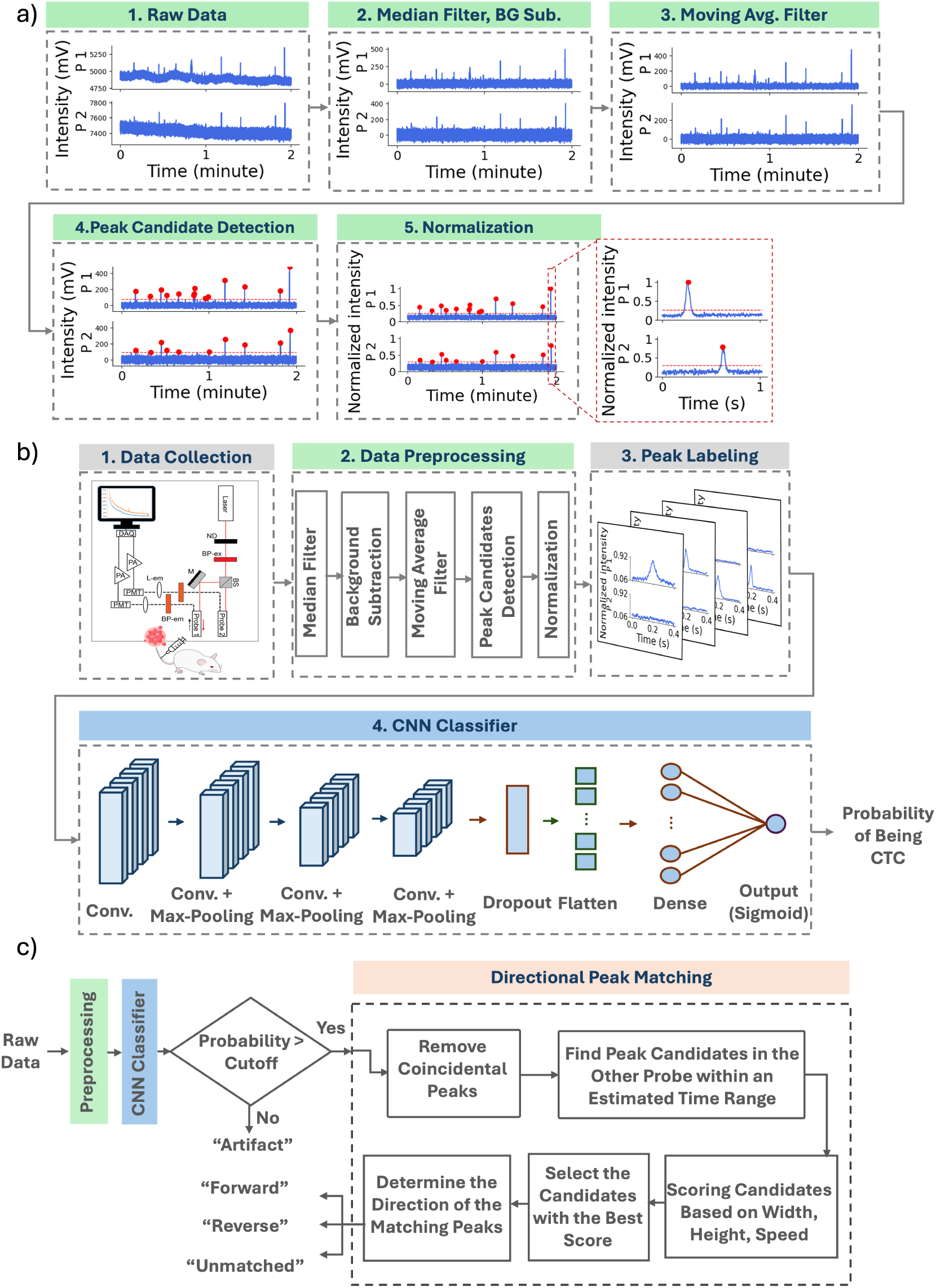
Overview of the ML-integrated signal processing. (a) Data pre-processing steps (BG Sub = background subtraction). (b) ML-based peak classification: After data collection, pre-processing, and labeling, the data were passed through the CNN for the training phase, yielding the probability of each peak candidate being a true peak. (c) Directional peak matching model determined the direction of CTC movement.

#### 2.2.2 CNN Classifier

The CNN classifier was designed with 4 two-dimensional convolutional layers, each with 8 filters (Fig. 3b). The kernel sizes for these layers were 1×64, 1×32, 1×16, and 1×8, respectively. With the exception of the first one, each convolutional layer was followed by a max-pooling layer with a kernel size of 1×4 to reduce the spatial dimensions of the output. Max-pooling was not applied after the first layer to avoid excessive down-sampling that could lead to invalid (negative) feature dimensions. To reduce overfitting, a dropout layer was introduced after the convolutional layers, randomly setting 25% of input units to zero during each training step.^23^ The resulting output was then flattened and passed to a fully connected neural network with two dense layers. The Rectified Linear Unit (ReLU) was used as the activation function for the convolutional layers and the first dense layer.^24^ The final dense layer employed a sigmoid activation function to produce an output between 0 and 1, representing the probability that the analyzed window contains a true peak arising from a fluorescently-labeled or fluorescence-expressing cell.

The CNN was trained on manually labeled windows (see Sec. 2.2.3) using the Adam optimizer and binary cross-entropy loss with 5-fold cross-validation. Training was conducted over 30 epochs, with early stopping applied if the validation loss did not improve for 3 consecutive epochs.^25^

The trained model was then applied to unseen DiFC data. Signal windows containing candidate peaks with predicted probabilities exceeding a predefined cutoff were classified as “1” (CNN-predicted peaks or Peak_CNN_) and passed to the directional matching algorithm for final classification, while all remaining windows were classified as “0” or artifacts. The probability cutoff was determined based on receiver operating characteristic (ROC) analysis and the distribution of predicted probabilities (see Sec. 3.1).

#### 2.2.3 Peak Labeling and CNN Training

The data sets used in this study were drawn from our previously published work as follows:

1. Multiple myeloma (MM) Disseminated Xenograft Model (DXM).^7^ Briefly, 8-week-old male SCID-bg mice were intravenously (i.v.) injected via the tail vein with MM.1S.GFP.Luc cells. After injection, MM cells rapidly homed to the bone marrow niche and proliferated, with CTCs beginning to disseminate into the bloodstream ∼3 weeks later. Two cohorts of mice were monitored for up to ∼5 weeks post-injection. A blue-green GFP–compatible DiFC system operating at 2000 Hz was used to scan the tail of the mice bi-weekly, with each scan lasting 45 min.
2. L1210A Leukemia Model,^8^.^9^ Briefly, female athymic nude mice received L1210A leukemia cells labeled with a near-infrared fluorescent contrast agent at 6-8 weeks of age (OTL38,^26^). L1210A cells were either labeled *in vitro* with OTL38 prior to i.v. injection, or L1210A cells and OTL38 were i.v. injected separately into the same mouse. The tail or leg was then scanned with a near-infrared–compatible DiFC system operating at 2000 Hz for 1 hour.

A total of 18 scans from CTC-bearing mice were used for manual labeling of the training and test datasets. These comprised 11 scans from the MM DXM (obtained from 7 SCID-bg mice) and 7 scans from the L1210A model (obtained from 3 athymic nude mice). In addition, 12 DiFC scans were acquired from 5 control mice injected with PBS only, including 2 SCID-bg mice scanned using the GFP–compatible DiFC system^7^ and 3 ahtymic nude mice scanned using the near-infrared– compatible DiFC system,^8^.^9^ Individual scans lasted from 45 min to 1 hour.

For training, peak candidates identified during pre-processing were manually inspected and labeled as “1” if attributed to a fluorescent cancer cell passing through the optical fiber, and as “0” if attributed to artifacts such as electrical noise, motion, or breathing. Labeling was performed by trained readers using established morphological and temporal criteria to distinguish true peaks from artifacts. True peaks exhibited a characteristic bell-shaped profile with measurable width and distinct peak area (Figs. 4a,b), whereas electrical noise appeared as abrupt, sharp spikes (Fig. 4c). In addition to peak shape, a matching peak detected in the signal from the other probe with an expected time delay was required to confirm a true positive. Conversely, coincident peaks appearing simultaneously in the signals from probes 1 and 2 were considered artifacts (Fig. 4d). Any peak candidates identified in scans performed on control mice were automatically labeled as artifacts since it was known *a priori* that no fluorescently-labeled cells were present in circulation. In total, 2,200 true peaks (”1”) and 4,099 artifacts (”0”) were annotated. To construct balanced training and test datasets, 2,200 artifacts were randomly selected from the full artifact set. Combined with the 2,200 true peaks, this yielded a dataset of 4,400 windows, 3,520 of which were randomly designated for the training set and the remaining 880 were used for the test set. This labeled training set enabled the network to learn distinguishing features between true peaks and artifacts based on their shape and other signal characteristics. The CNN classifier was trained on an Apple M3 Pro processor with 18 GB of memory and required ∼31 min of total CPU time.

**Fig 4.**
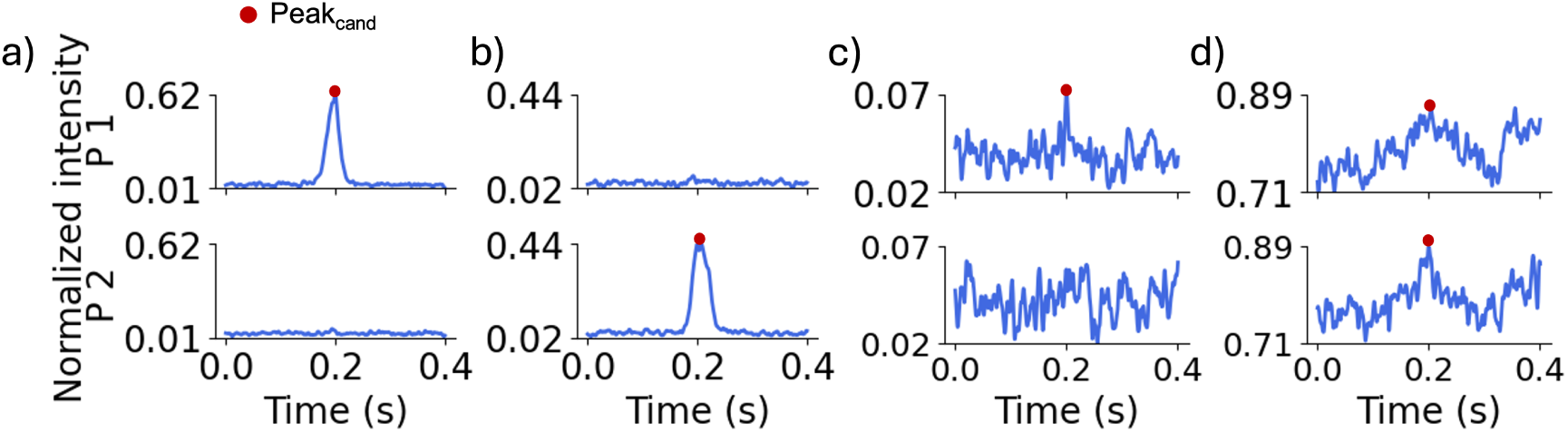
Example 0.4 s data windows centered on peak candidates showing, a true peak from a fluorescently-labeled cell detected by (b) probe 1 and (c) probe 2, respectively. False peak candidates may originate from (c) electrical noise and (d) motion artifacts.

#### 2.2.4 Data Augmentation

Since manual data labeling was time-consuming, we used data augmentation to expand the training dataset.^27^ We applied two strategies: “flipped-window” and “swapped-probes”.^28^ In the flipped-window approach, each training example was reversed in time while retaining its original label. Figures. 5a and 5b show a representative true peak and its time-reversed counterpart, respectively.

**Fig 5.**
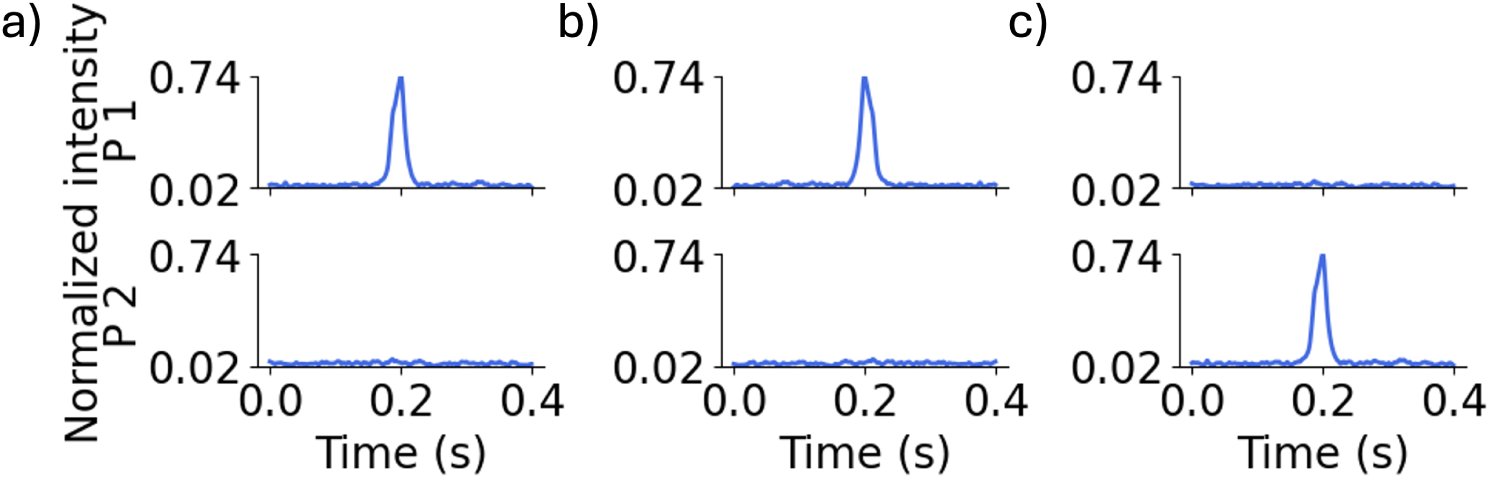
(a) Representative 0.4 s DiFC data window showing (a) a true peak in probe 1, (b) a flipped-window version of (a), and (c) a swapped-probe version of the window in (a).

In the swapped-probes approach, the signals from probes 1 and 2 were exchanged, such that a true peak originally presents in probe 1 appeared in probe 2 after swapping, and vice versa (Fig. 5c). Both data augmentation methods were applied to a randomly-selected 25% subset of the training set, expanding the training set from 3,520 to 5,280 examples.

### 2.3 Directional Peak Matching

The directional peak matching algorithm was applied as previously described.^6, 7^ Briefly, coincident peaks appearing simultaneously on both probes were excluded. For each remaining peak, a search interval was defined using an estimated cell speed derived from peak width. Potential matches within this interval were identified, and the best match was selected based on similarity in peak width, height, and inferred speed. Each matched pair was then labeled as “forward” or “reverse” based on relative peak timing, and candidates without a suitable match were recorded as “unmatched”. In the ML-integrated signal processing approach, only peak candidates with a CNN-predicted probability indicating a true peak (Peak_CNN_), rather than an artifact, were passed to the directional peak-matching algorithm.

### 2.4 Validation Data Sets

To validate the proposed algorithm, evaluate its performance on unseen data, and compare it with the threshold-based method, we used DiFC scans from *in silico* datasets, control mice, and CTC-bearing mice, as follows.

#### 2.4.1 In Silico Data

Prior to testing our algorithm on experimental DiFC data, we generated an *in silico* dataset to test model performances under known conditions. In the *in silico* scan, the ground truth signal (i.e., the exact number, type, direction, and timing of detections) was known. We generated a library of three building block types: background noise, true peak, and artifact blocks, each extracted from real *in vivo* data that were excluded from all training, testing, and evaluation sets. True peak- and artifact- containing blocks consisted of 0.4 s windows centered on manually labeled peak candidates, while background noise blocks contained pre-processed autofluorescence signal with minimal fluctuations. Blocks were selected with near zero endpoint values and small variation to ensure smooth stitching and were then concatenated to generate a 10 min *in silico* scan. We used a structured sequence of segments: background noise, artifact in probe 2, true peak in probe 1, true peak in probe 2, artifact in probe 1, and background noise again (Fig. 6), where co-occurring true peaks and artifacts may lead to incorrect directional matches. This sequence was repeated 40 times, with background noise blocks inserted between instances to generate complete *in silico* data.

**Fig 6.**
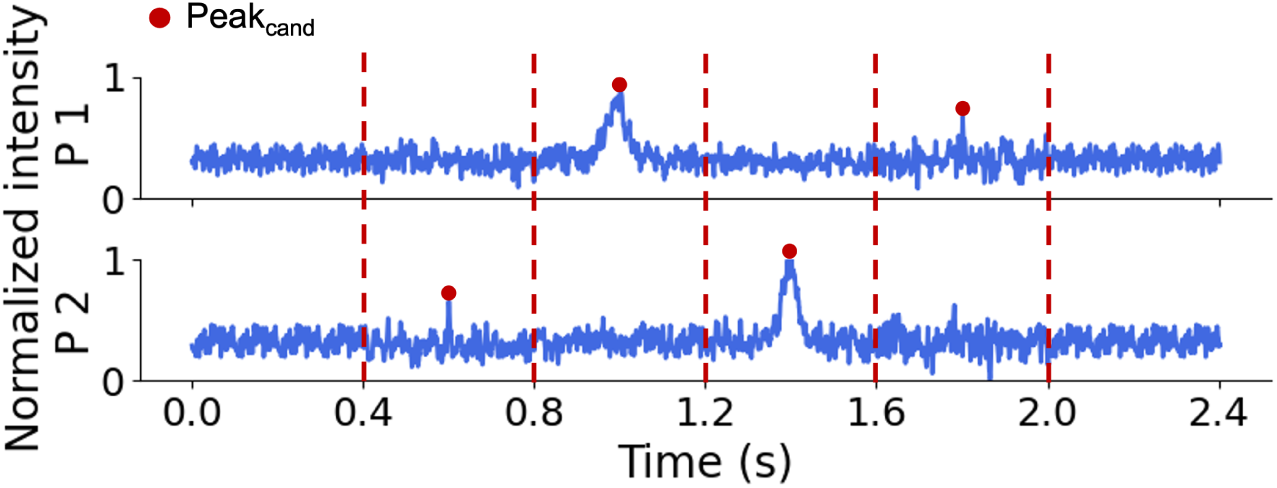
Building block sequence used to generate the *in silico* dataset. From left to right: background noise, artifact in probe 2, true peak in probe 1, true peak in probe 2, artifact in probe 1, and background noise.

#### 2.4.2 Control (Non-CTC Bearing) Mice

Scans from control, non-CTC bearing mice (Sec. 2.2.3) were used to evaluate algorithm performance (since all detected peak candidates were, by definition, artifactual). We used a total of 6 scans lasting 45 min each: 3 acquired with the GFP–compatible DiFC system and 3 with the near-infrared–compatible DiFC system.

#### 2.4.3 CTC-Bearing Mice

We analyzed 360 min of data from 8 scans acquired in CTC-bearing mice, half of which from GFP-expressing MM-bearing mice and half from mice injected with labeled L1210A cells (Sec. 2.2.3). Unlike for the *in silico* and control data sets, ground truth was not known *a priori*, therefore each scan was manually reviewed and labeled.

### 2.5 Metrics

To evaluate candidate peak classification performance (Peak_CNN_ vs. artifacts) of the CNN classifier, we computed accuracy, precision, sensitivity, specificity, as defined in Eqs. (1)–(4), using the test dataset described in Sec. 2.2.3.

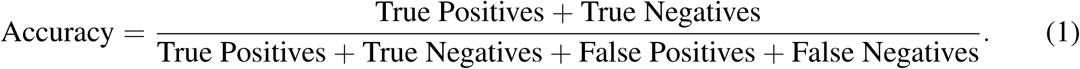

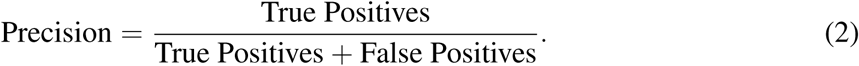

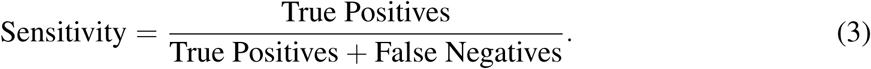

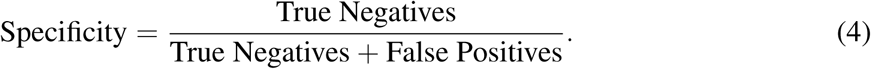

We also calculated the the area under the ROC curve (AUC), which illustrated the relationship between True Positive Rate (TPR) and False Positive Rate (FPR) across classification probability cutoff values. The TPR measured the proportion of true peaks correctly identified by the model and was equivalent to sensitivity, as defined in Eq. (3). The FPR quantified the proportion of artifacts incorrectly classified as true peaks and was computed as the ratio between the number of false positives and the total number of artifacts, defined in Eq. (5).

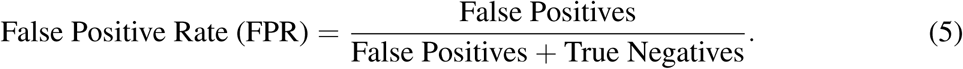

To evaluate the directional peak matching algorithm, we assessed whether matches were correctly or incorrectly classified by direction. Correct matches were true forward or reverse peak pairs correctly identified by the algorithm, whereas incorrect matches were artifact or unmatched peaks misclassified as forward or reverse. For clarity in the evaluation of the ML-integrated approach, key classification terms are defined in Table 1.

**Table 1.** Description of key classification terms used in ML-integrated DiFC signal processing. F, R and U represents “forward”, “reverse” and “unmatched”, respectively.

| Term | Brief description |
| --- | --- |
| Peak (true peak) | Transient fluorescent signal caused by a CTC detected in the FOV. |
| Artifact | Signal fluctuation from non-biological sources (e.g. noise, motion). |
| Peak<sub>cand</sub> | Signal transient exceeding threshold; potential peak before validation. |
| Peak<sub>CNN</sub> | Peak candidate classified by the CNN as a peak. |
| Forward | Peak pair indicating CTC movement from first to second probe. |
| Reverse | Peak pair indicating CTC movement from second to first probe. |
| Unmatched | Peak with no match on the alternate probe in the search interval. |
| Missed | Peak below detection threshold, excluded as a peak candidate. |
| True positive | Peak correctly classified by CNN or labeled F/R/U in directional matching. |
| False positive | Artifact incorrectly accepted by CNN or labeled F/R/U in directional matching. |
| True negative | Artifact correctly rejected by CNN or directional matching. |
| False negative | Peak incorrectly rejected by CNN or directional matching. |
| Correct match | Correctly identified forward or reverse peak pairs. |
| Incorrect match | Artifacts or unmatched peaks incorrectly labeled as pairs. |

## 3 Results

### 3.1 Model Training and Characterization

We first characterized the performance of the CNN classifier without the use of directional matching and determined the binary cross-entropy loss over epochs for both the training and validation sets (Fig. 7a). Early stopping was applied during training, based on validation loss, with a patience of 3 epochs. This setting restored the model to its best weights before training is stopped. Due to the use of regularization and dropout layers during the training phase to enhance generalization, the training loss was occasionally slightly higher than the validation loss. As illustrated in this figure, the training stopped at the 19^th^ epoch.

**Fig 7.**
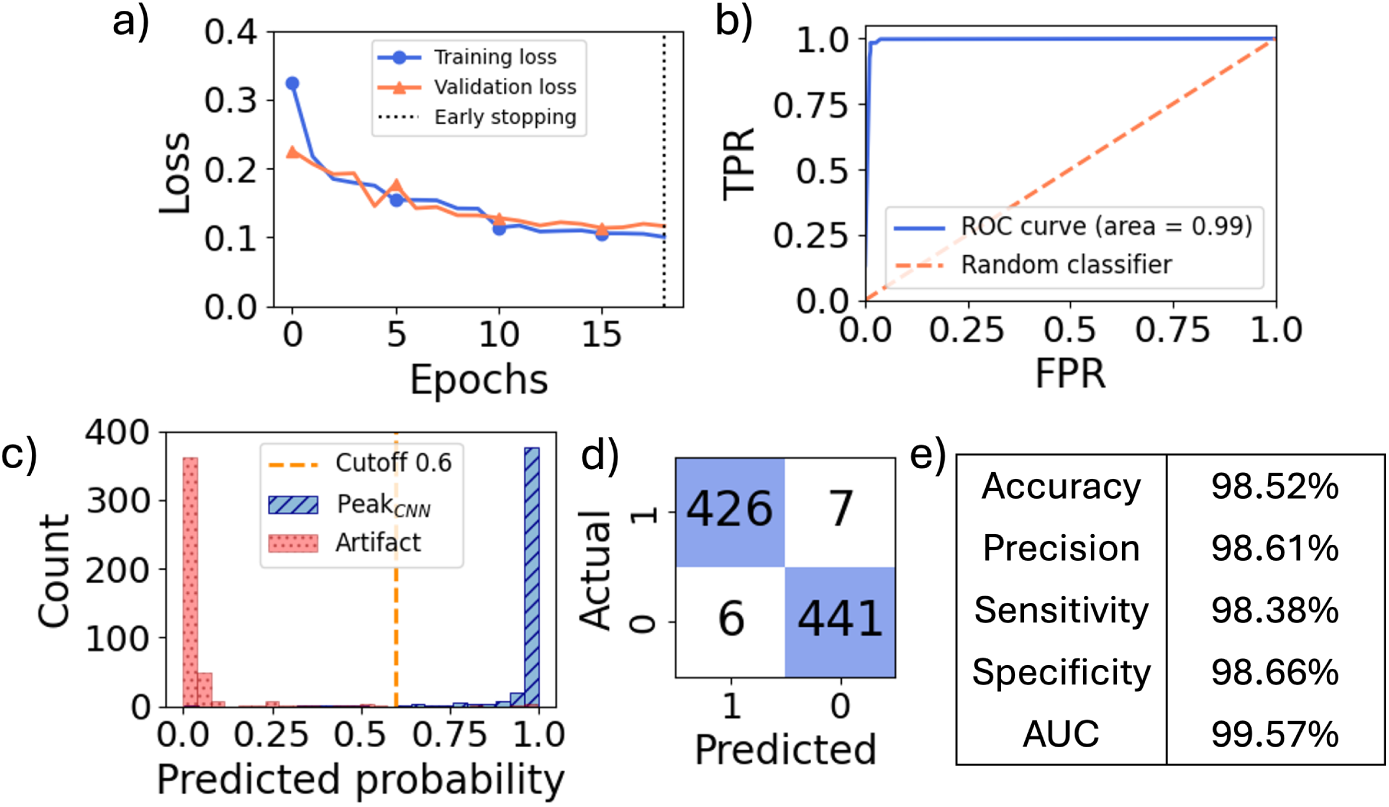
(a) Training loss and validation loss over training epochs. Training stopped at the 19^th^ epoch when the validation loss did not improve for three consecutive epochs. (b) ROC curve for the CNN classifier. (c) Histogram of the predicted probabilities for peak candidates in the test set, with true peaks shown in blue and artifacts shown in red. As shown, predicted probabilities for true peaks are close to 1, while those for artifacts are close to 0. The orange dashed line indicates the selected probability cutoff for the CNN classifier. (d) Peak classification confusion matrix using the CNN classifier on the test set with the cutoff set to 0.6. “0” and “1” represent artifact and peak, respectively. (e) Accuracy, precision, sensitivity, specificity, and AUC of the CNN classifier on the test data set.

We next evaluated the trained CNN model on the test dataset. The resulting ROC curve generated by varying the classifier probability cutoff in the range of [0 to 1] is shown in Fig. 7b.

The distribution of predicted probabilities for both true peaks and artifacts in the test set was also analyzed (Fig. 7c). As shown, the predicted probabilities for true peaks (blue) were close to 1, while those for artifacts (red) were near 0. By examining both the ROC curve and the predicted probability distribution, we selected 0.6 as a peak-candidate cutoff that balanced low-false positive and high-true positive performance, although this can be adjusted to trade off the two considerations. This cutoff resulted in 7 false positives and 6 false negatives while retaining high true positives (426) and true negative (441) counts. We determined the accuracy (98.52%), precision (98.61%), sensitivity (98.38%), and specificity (98.66%) of the model, as shown in the corresponding confusion matrix and performance metrics for the test set (Figs. 7d,e).

### 3.2 Performance on DiFC Data Sets

#### 3.2.1 In-silico Data

In general, the ground truth number and direction of true peaks is not known for any DiFC dataset measured in CTC-bearing mice *in vivo*. To this end, we synthetically generated a challenging 10 min scan containing 80 artifacts and 80 true peaks (all of which were in the forward direction) as described in Sec. 2.4.1. The results are summarized in Fig. 8, where the first and middle row illustrate the result from our previously documented threshold-based approach^6, 7^ with the threshold of 5*σ*_bg_, and 4*σ*_bg_, respectively. the last row corresponds to the results from the ML-integrated approach with the threshold of 4*σ*_bg_.

**Fig 8.**
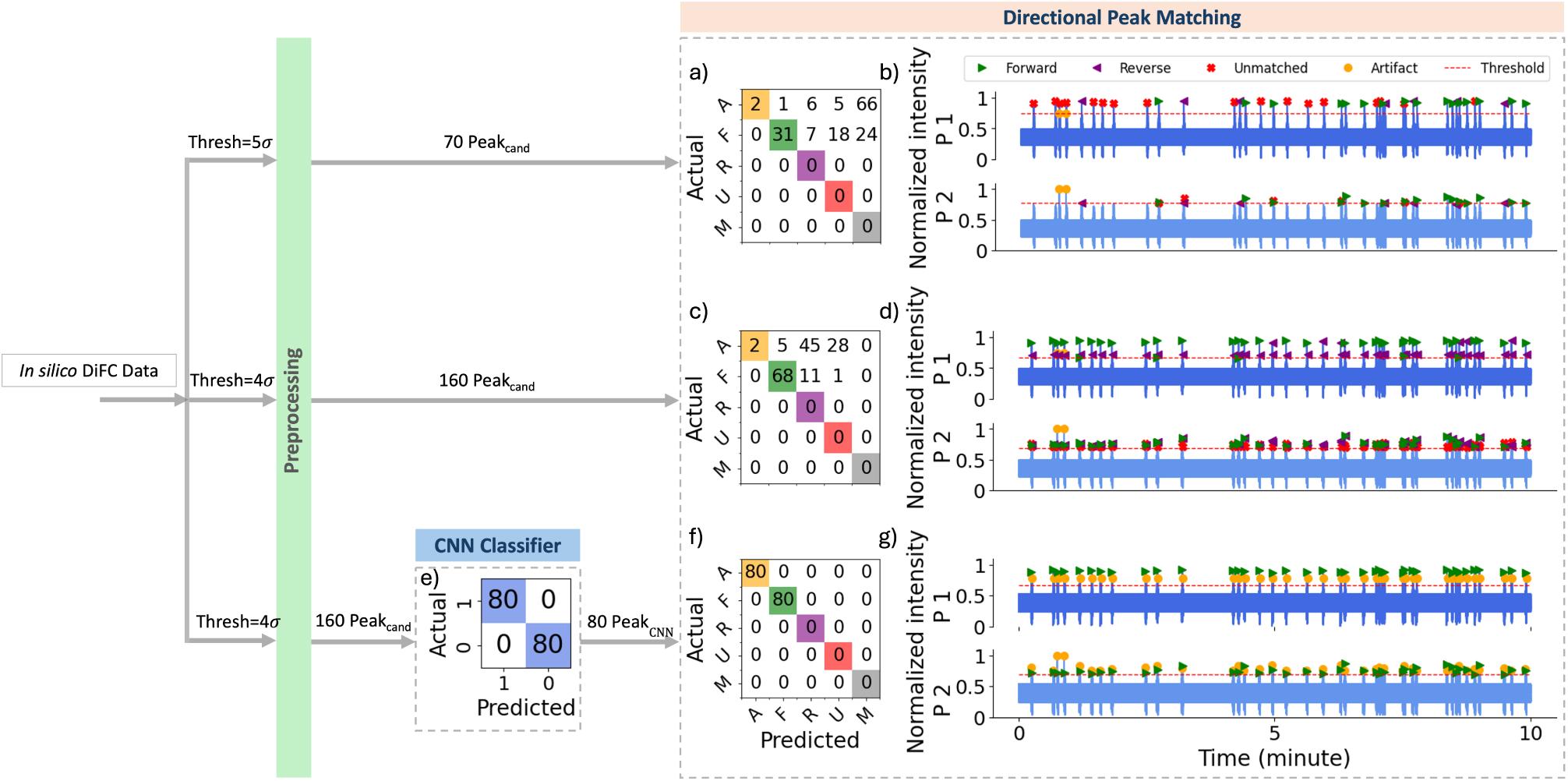
Performance comparison of the directional peak matching models on 10 min of *in silico* data. (a) Confusion matrix for the threshold-based directional peak matching with a threshold of 5*σ*_bg_. “A,” “F,” “R,” “U,” and “M” are the abbreviated forms of artifact, forward, reverse, unmatched, and missed, respectively. (b) Color-coded predicted labels using the same model in (a). (c) Confusion matrix for the threshold-based directional peak matching with a threshold of 4*σ*_bg_. (d) Color-coded predicted labels using the same model in (c). (e) Confusion matrix for the CNN classification on peak candidates detected with the threshold of 4*σ*_bg_. “0” and “1” represent artifact and peak, respectively. (f) Confusion matrix for the ML-integrated directional peak matching with a threshold of 4*σ*_bg_. (g) Color-coded predicted labels using the same model in (f).

The threshold-based approach with a threshold of 5*σ*_bg_ identified 70 peak candidates for further analysis with the directional peak matching algorithm, whereas a further 90 lower amplitude peaks were missed as shown in the last column of the extended confusion matrix in Fig. 8a. This resulted in fewer true forward or reverse CTCs compared to using a lower threshold of 4*σ*_bg_, though the higher threshold also reduced false positives by avoiding dimmer artifacts, as demonstrated in Fig. 8a.

Lowering the threshold to 4*σ*_bg_ (Figs. 8c-d) allowed us to capture more peak candidates as is evident by the increase in counts in the upper-left quadrant of Fig. 8c, and by the elimination of peaks in the missed column in Fig. 8c. This resulted in an increase of the correctly categorized as forward direction peaks (from 31 in Fig. 8a to 68 in Fig. 8c). However, this was accompanied by a rise in the number of false positives, 45 and 5 of which were erroneously labeled as reverse and forward directions, respectively.

Thus, while lowering the threshold increased the detection of true positives and improved the matching of forward peaks, it also led to a considerable increase in false positives and incorrect matches, illustrating the trade-off between sensitivity and specificity in the threshold-based approach.

Next, with the lower threshold of 4*σ*_bg_, we applied our CNN classifier prior to the peak matching algorithm (”ML-integrated approach”). As illustrated in the peak classification confusion matrix, Fig. 8e, the CNN classifier correctly classified all true peaks and artifacts. Only peak candidates classified as “1” (Peak_CNN_) were passed to the directional peak matching step, removing artifacts from further analysis in the pipeline.

Following the directional matching step, all Peak_CNN_ were correctly labeled as forward, as shown in Fig. 8f. This demonstrates that the ML-integrated approach not only increased true positives and correct matches but also maintained a low number of false positives and incorrect matches.

#### 3.2.2 Non-CTC Bearing Control Mouse

We next tested our ML-integrated approach on DiFC data sets measured non-CTC bearing mice. Analysis of a typical control (non-CTC bearing mouse) data set is shown in Fig. 9. As shown in Fig. 9a, application of our previously reported threshold-based method with a 5*σ*_bg_ threshold resulted in detection of no peak candidates (and hence no false positive detections). However, when the threshold was lowered to 4*σ*_bg_ (Fig. 9c), 72 artifacts were identified as peak candidates, 4 of which were misclassified as forward and 68 as unmatched by the directional matching algorithm, respectively.

**Fig 9.**
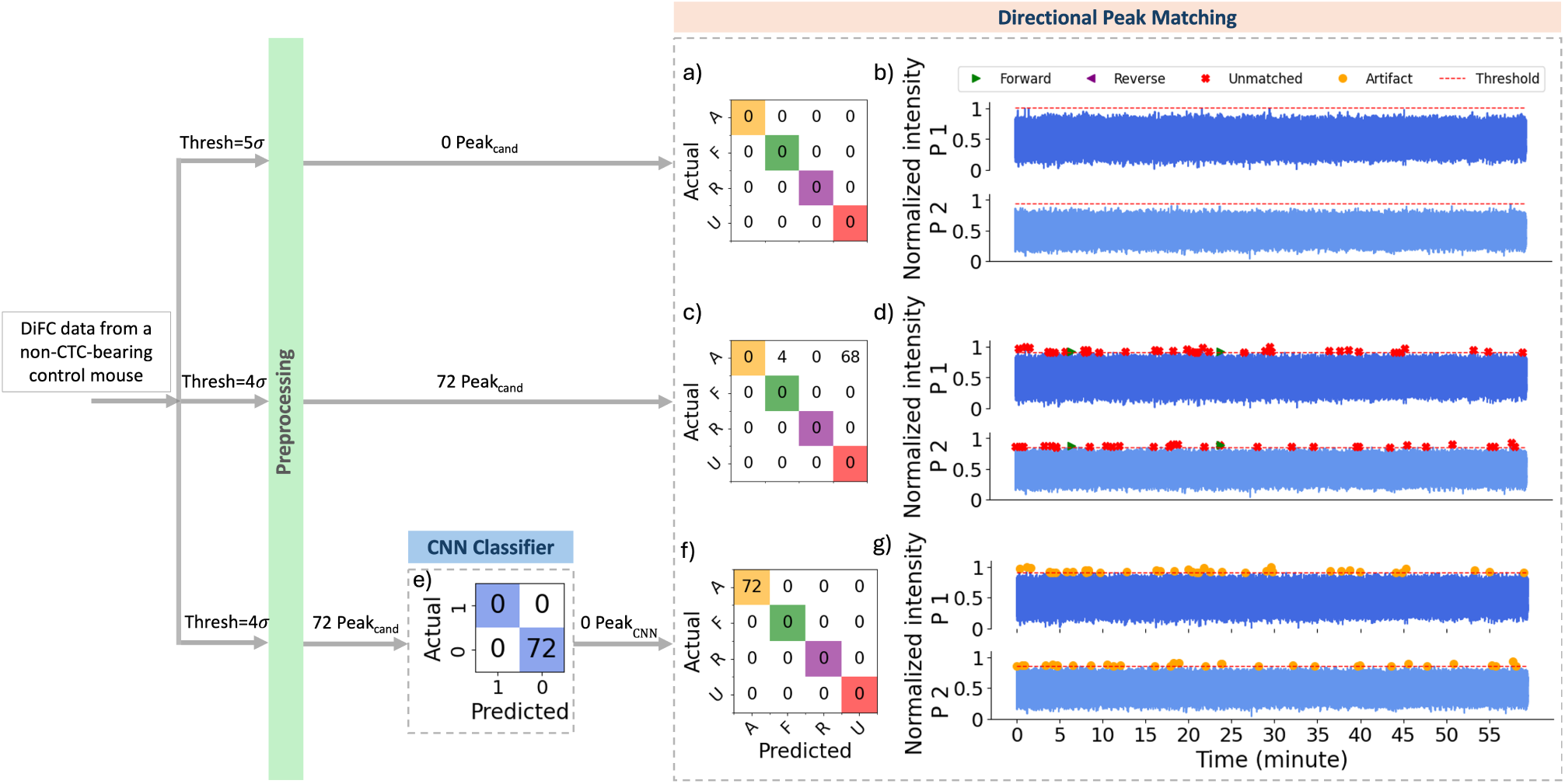
Performance comparison of the threshold-based and ML-integrated directional peak matching model on 45 min of scan collected from control mouse. (a) Confusion matrix for the threshold-based directional peak matching with a threshold of 5*σ*_bg_. (b) Color-coded predicted labels using the same model in (a). (c) Confusion matrix for the threshold-based directional peak matching with a threshold of 4*σ*_bg_. (d) Color-coded predicted labels using the same model in (c). (e) Confusion matrix for the CNN classification on peak candidates detected with the threshold of 4*σ*_bg_. (f) Confusion matrix for the ML-integrated directional peak matching with a threshold of 4*σ*_bg_. (g) Color-coded predicted labels using the same model in (f).

With the application of the CNN classifier after pre-processing (Fig. 9e), all 72 artifacts captured as peak candidates were classified as “0” (artifact), and none proceeded to directional matching, resulting in no false positives, as illustrated in Fig. 9f.

Analysis of a particularly noisy individual control dataset is shown in Fig. 10. Application of the threshold-based approach with a threshold of 5*σ*_bg_ resulted in detection of 42 peak candidates, 32 of which were classified as unmatched by the directional matching algorithm and 10 as artifact through coincident peak removal.

**Fig 10.**
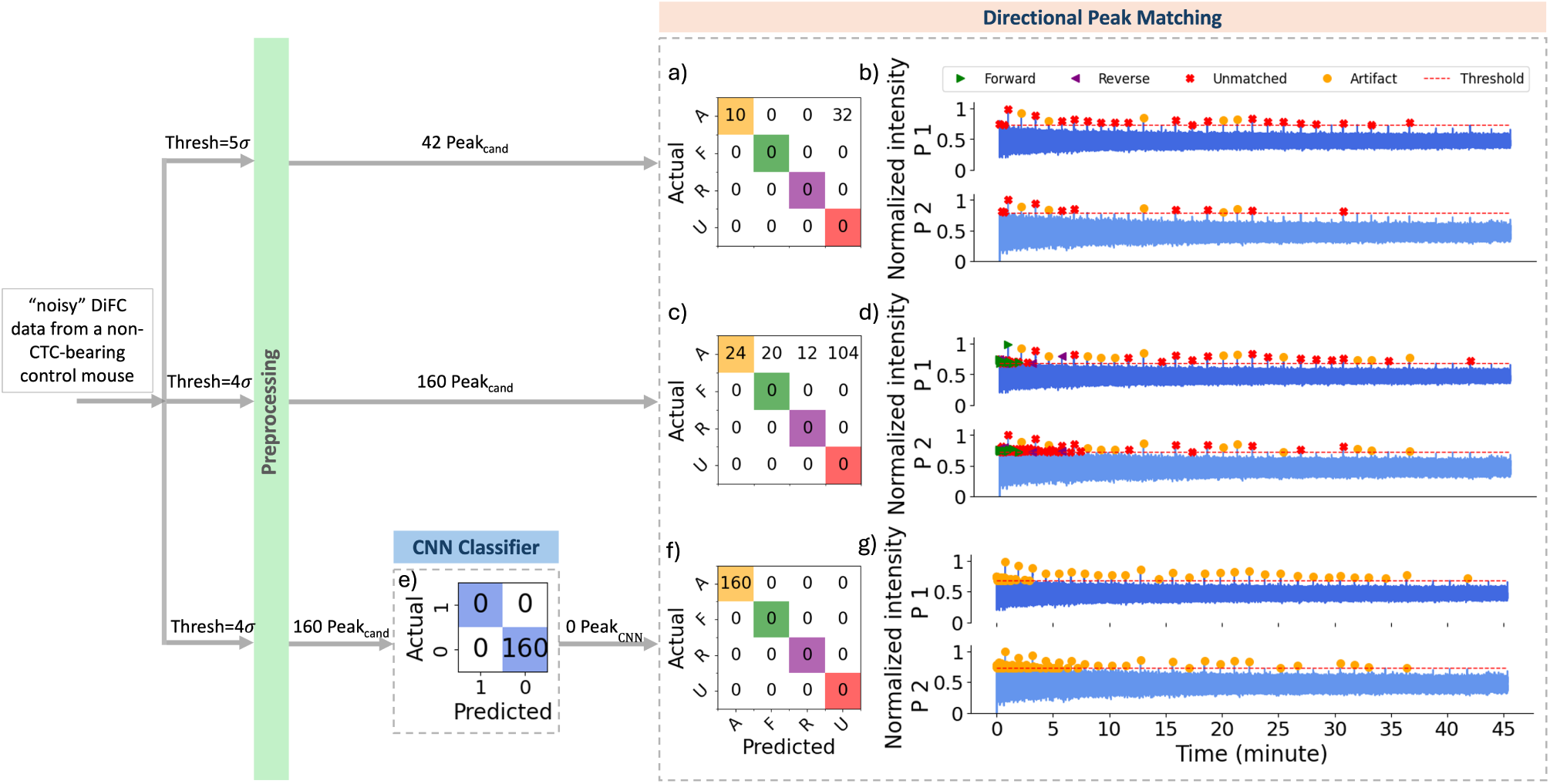
Performance comparison of the threshold-based and ML-integrated directional peak matching model on 45 min of noisy scan collected from control mouse. (a) Confusion matrix for the threshold-based directional peak matching with a threshold of 5*σ*_bg_. (b) Color-coded predicted labels using the same model in (a). (c) Confusion matrix for the threshold-based directional peak matching with a threshold of 4*σ*_bg_. (d) Color-coded predicted labels using the same model in (c). (e) Confusion matrix for the CNN classification on peak candidates detected with the threshold of 4*σ*_bg_. (f) Confusion matrix for the ML-integrated directional peak matching with a threshold of 4*σ*_bg_. (g) Color-coded predicted labels using the same model in (f).

When the threshold was lowered to 4*σ*_bg_, 160 peak candidates were detected, of which 128 were classified as unmatched or coincident peaks. Of the remaining peaks, 20 and 12 were erroneously matched in forward and reverse directions, respectively (Fig. 10c). In contrast, using the same threshold of 4*σ*_bg_, the CNN model accurately classified all peak candidates as “0” (artifacts), resulting in no erroneous directional peak matches (as shown in Fig. 10f). This demonstrated the robustness of the ML-integrated approach in rejecting artifacts even under noisy conditions.

#### 3.2.3 CTC-Bearing Mice

Analysis of a DiFC scan measured from an MM-DXM bearing mouse is summarized in Fig. 11. Application of the threshold-based method with a threshold of 5*σ*_bg_ resulted in detection of 46 peak candidates, of which only 22 were correctly forward-matched and 14 were correctly unmatched peaks. Lowering the threshold to 4*σ*_bg_ increased the number of detected candidates to 119; using the threshold-based method, this yielded 42 correct forward matches and 26 correct unmatched peaks, however at the cost of significantly increased false positives (40 artifacts labeled as forward, reverse, or unmatched) and incorrect matches (6 artifacts matched as forward and 5 as reverse) after directional peak matching (Fig. 11b). Using the ML-integrated approach with a threshold of 4*σ*_bg_, all 119 detected peak candidates were correctly classified by the CNN classifier as 45 artifacts and 74 peaks (Fig. 11e). Subsequent directional matching of the 74 peaks yielded 42 correct forward matches and 26 correct unmatched peaks (Fig. 11f).

**Fig 11.**
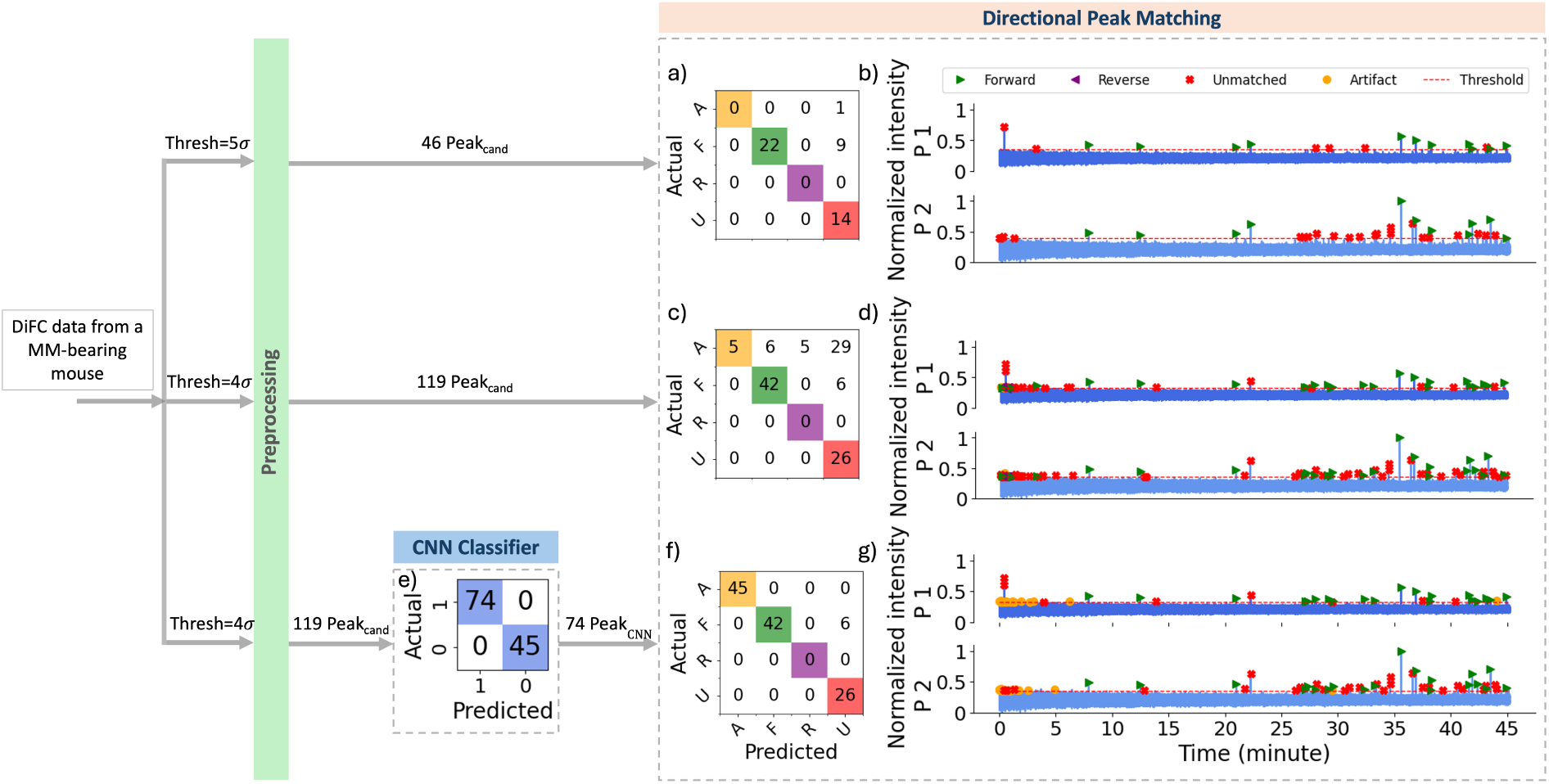
Performance comparison of the directional peak matching models on 45 min of data collected from MM-bearing mouse. (a) Confusion matrix for the threshold-based directional peak matching with a threshold of 5*σ*_bg_. (b) Color-coded predicted labels using the same model in (a). (c) Confusion matrix for the threshold-based directional peak matching with a threshold of 4*σ*_bg_. (d) Color-coded predicted labels using the same model in (c). (e) Confusion matrix for the CNN classification on peak candidates detected with the threshold of 4*σ*_bg_. (f) Confusion matrix for the ML-integrated directional peak matching with a threshold of 4*σ*_bg_. (g) Color-coded predicted labels using the same model in (f).

These results highlight the ML-integrated model’s ability to address the limitations of threshold-based methods across varying thresholds.

We next analyzed data in the same manner from L1210A-bearing mice. Lowering the threshold from 5*σ*_bg_ to 4*σ*_bg_ in the threshold-based approach increased the number of peak candidates from 145 to 169, and consequently increased the number of false positives to 15 (15 artifacts labeled as unmatched), as shown in Figs. 12a,c.

**Fig 12.**
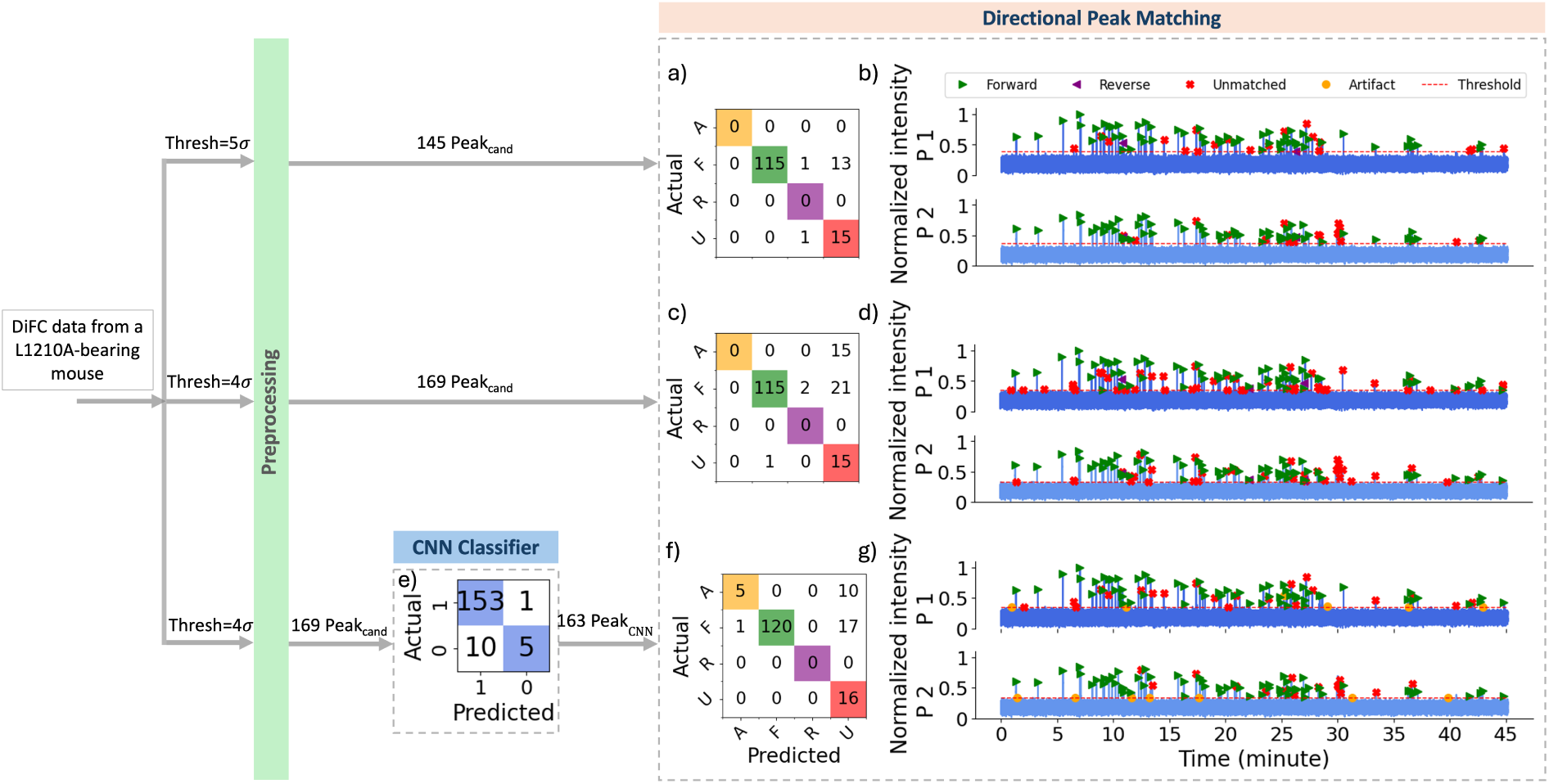
Performance comparison of the directional peak matching models on 45 min of data collected from L1210A-bearing mouse. (a) Confusion matrix for the threshold-based directional peak matching with a threshold of 5*σ*_bg_. (b) Color-coded predicted labels using the same model in (a). (c) Confusion matrix for the threshold-based directional peak matching with a threshold of 4*σ*_bg_. (d) Color-coded predicted labels using the same model in (c). (e) Confusion matrix for the CNN classification on peak candidates detected with the threshold of 4*σ*_bg_. (f) Confusion matrix for the ML-integrated directional peak matching with a threshold of 4*σ*_bg_. (g) Color-coded predicted labels using the same model in (f).

In the ML-integrated approach with a threshold of 4*σ*_bg_, the 169 peak candidates were first processed by the CNN classifier, which filtered out 6 candidates (represented in the “0” column of the 2 × 2 confusion matrix in Fig. 12e, including 5 actual artifacts). The remaining 163 peak candidates were then passed to the directional peak matching algorithm. This reduced false positives from 15 to 10 and increased the number of correct forward matches from 115 to 120 compared to the threshold-based method, with no incorrect matches after directional peak matching (Fig. 12f). We note that the extended confusion matrices shown in the output of directional peak matching for the *in vivo* validation datasets (Figs. 9, 10, 11, 12) do not include a “missed” row or column, unlike the matrices for *in silico* scans in Fig. 8, i.e. because the exact number of true peaks was not known a priori.

#### 3.2.4 Overall Results

Figure. 13 summarizes the performance of the algorithm for all control and CTC-bearing data used in this study using the threshold-based and ML-integrated methodology described in Sec. 2.4.2 and Sec. 2.4.3. As shown in Figs. 13a-c, lowering the threshold in the threshold-based method significantly increased the number of false positives from 55 to 873 whereas the ML-integrated approach produced no false positives, with all 903 artifacts in control scans correctly classified. For the CTC-bearing scans, although lowering the threshold to 4*σ*_bg_ increased the number of correct matches (from 400 correct forward and 4 correct reverse peaks to 602 correct forward and 9 correct reverse peaks; Figs. 13d,e), it also significantly increased false positives (from 11 to 373) and incorrect matches (from 17 to 66). The ML-integrated approach at the same threshold not only yielded a comparable number of correct matches (607 correct forward and 10 correct reverse peaks) but also maintained substantially fewer false positives (23) and incorrect matches (25), as illustrated in Fig. 13f.

**Fig 13.**
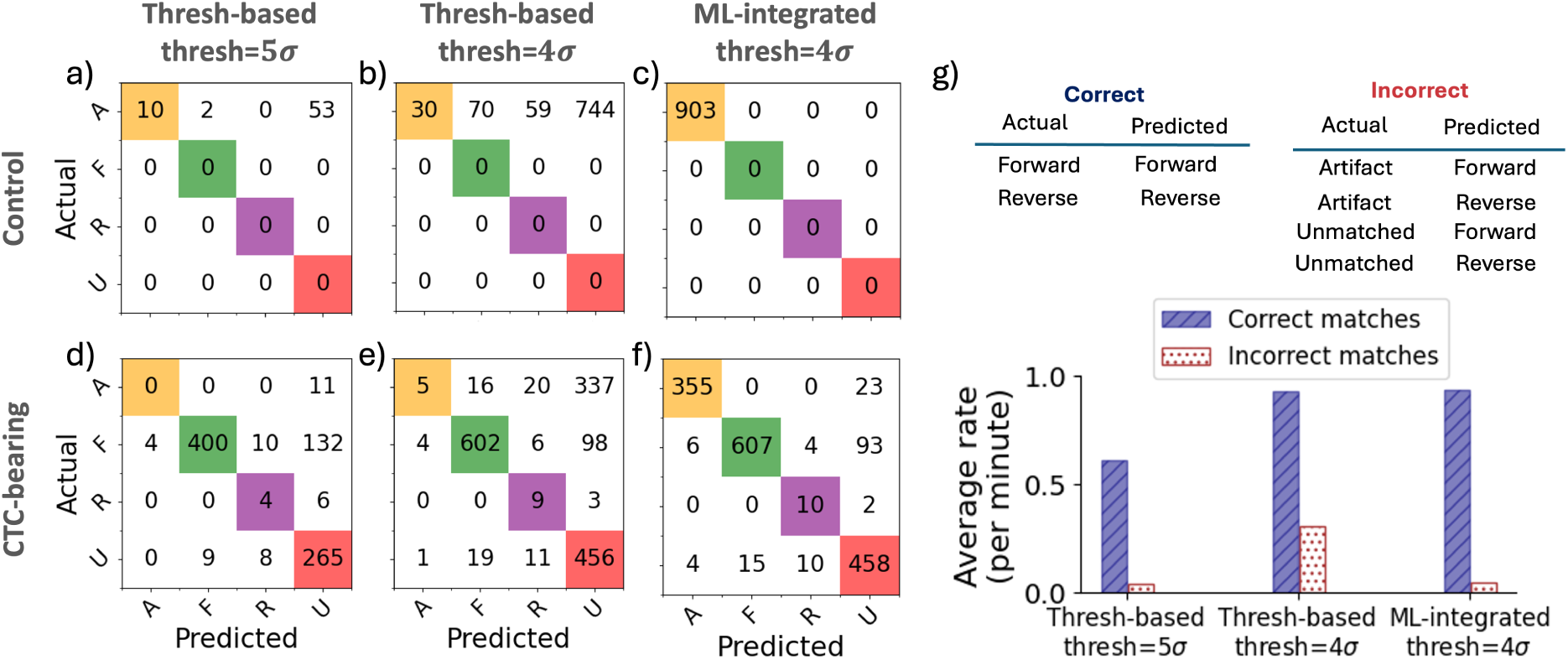
Overall performance of threshold-based and ML-based directional peak matching, derived from manually labeled validation scans collected from control and CTC-bearing mice. (a), (b) Confusion matrix for threshold-based directional peak matching on control data with threshold of 5*σ*_bg_ and 4*σ*_bg_, respectively. (c) Confusion matrix for ML-integrated directional peak matching on control data with threshold of 4*σ*_bg_. (d),(e) Confusion matrix for threshold-based directional peak matching on CTC-bearing mice data with threshold of 5*σ*_bg_ and 4*σ*_bg_, respectively. f) Confusion matrix for ML-integrated directional peak matching on CTC-bearing mice data with threshold of 4*σ*_bg_. (g) Bar plot of correct matches vs. incorrect matches. Correct matches are defined as forward or reverse peaks correctly labeled. Incorrect matches are defined as artifact or unmatched peaks incorrectly labeled as forward or reverse.

Last, Fig. 13g shows the correct versus incorrect matches for the threshold-based approach with thresholds of 5*σ*_bg_ and 4*σ*_bg_, and for the ML-integrated approach with a threshold of 4*σ*_bg_. These results are based on all validation scans from both control and CTC-bearing mice, as described in Sec. 2.4.2 and Sec. 2.4.3. As shown, the ML-integrated approach achieved a high number of correct matches (comparable to the threshold-based approach at 4*σ*_bg_) while maintaining a low number of incorrect matches (comparable to the threshold-based approach at 5*σ*_bg_), effectively addressing the limitations of the threshold-based models.

## 4 Discussion

In this study, we developed and validated a CNN classifier for peak classification in DiFC and demonstrated its robustness with data acquired using our blue-green (GFP–compatible) and near-infrared–compatible DiFC systems, and using two different cell lines and mouse strains. By incorporating scans collected from different cancer cell lines and using both GFP–compatible and near-infrared–compatible DiFC systems in the training, testing, and validation datasets, we ensured the model’s adaptability and independence from specific experimental setups. This versatility highlights the potential of the CNN classification as a generalized solution for DiFC applications.

The results consistently showed that the CNN classifier addresses key limitations of threshold-based methods. Specifically, when using a high threshold, threshold-based approaches reduce false positives but fail to capture lower SNR peaks, leading to missed true positives. Conversely, lowering the threshold captures more true positives but significantly increases false positives, especially in noisy datasets. The CNN classifier, however, incorporates both the shape as well as the amplitude of peak candidates, thereby achieving a balance by capturing a higher number of true positives while maintaining a low rate of false positives.

The advantages of the CNN classifier become particularly evident in scenarios with noisy data, where artifacts and fluctuations in the signal are prevalent. Threshold-based methods struggle in such conditions, often capturing artifacts as peak candidates and producing numerous false positives. In contrast, the CNN classifier effectively distinguishes between true peaks and artifacts, demonstrating its superior performance in challenging environments. When the SNR is high, and the data is relatively clean, threshold-based approaches can perform adequately; however, their utility diminishes in noisy conditions.

We next plan to deploy this approach in DiFC scans taken from awake, freely moving mice.^29^ In these scenarios, motion artifacts introduce significant challenges to the accuracy of signal interpretation. Threshold-based methods are prone to failure in such conditions, producing excessive false positives and compromising their reliability. The CNN classifier, with its ability to handle complex, noisy datasets, is well-suited to address these challenges, reinforcing its importance in the ongoing development of DiFC for dynamic and real-world experimental settings.

Likewise, this ML-enhanced approach to time-series signal processing may be useful for in vivo flow cytometry based approaches using intravital microscopy or photoacoustic designs.^3, 5, 30, 31^

## 4.2 Code, Data, and Materials Availability

The datasets analyzed during this work and the Python code are available through the Synapse repository (DOI: 10.7303/SYN73915900).

## 4.3 Acknowledgments

This work was supported by the National Institutes of Health/National Cancer Institute (NIH/NCI) under grant number 1R01CA260202.

We thank Dr. Qianqian Fang (Northeastern University) for valuable feedback on the manuscript.

**Mehrnoosh Emamifar** received her B.S. degree in Computer Engineering (2018) and M.S. degree in Artificial Intelligence (2021) from the University of Tehran. She is currently a Ph.D. candidate in Bioengineering at Northeastern University.

**Jane Lee** received her Bachelors (BS) in Biology from Wake Forest University and her Master of Liberal Arts (ALM) in the field of Biotechnology at the Harvard Extension School. She is currently a PhD Candidate in Bioengineering at Northeastern University.

**Joshua Pace** was a graduate student in the Bioengineering Department at Northeastern University at the time this research was conducted.

**Chiara Bellini** received her PhD in Biomedical Engineering from the University of Calgary in 2012. She is an Associate Professor and Associate Chair for PhD Program and Research of Bioengineering at Northeastern University.

**Mark Niedre** received his PhD from the University of Toronto in the Department of Medical Physics in 2004. He is a Professor of Bioengineering and Associate Dean for PhD Education at Northeastern University and a Fellow of SPIE.

